# Floral scent emission of *Epiphyllum oxypetalum*: identification of a novel cytosol-localized geraniol biosynthesis pathway

**DOI:** 10.1101/2023.11.26.568706

**Authors:** Yiyang Zhang, Yuhan Zhang, Qiurui Tian, Likun Wei, Ting Zhu, Zhiwei Zhou, Jiaqi Wang, Zhibin Liu, Wei Tang, Haijun Xiao, Mingchun Liu, Tao Li, Qun Sun

**Affiliations:** Key Laboratory of Bio-resources and Eco-environment Ministry of the Education, College of Life Sciences, Sichuan University, Chengdu, Sichuan 610064, PR China; Sichuan Academy of Botanical Engineering, Sichuan Academy of Agricultural Sciences, Zizhong, Sichuan 641200, PR China

**Keywords:** floral scent, VOCs, emission dynamics, transcriptome, terpene biosynthesis, MVA pathway

## Abstract

- *Epiphyllum oxypetalum*, a renowned ornamental species in Cactaceae, releases attractive fragrance during its infrequent, transient and nocturnal flowering, the nature and biosynthesis of the volatiles for this floral scent, however, remained unexplored.
- Employing volatilomic, transcriptomic and biochemical approaches, we systematically characterized the composition, emission dynamics and biosynthesis of *E. oxypetalum* floral scent.
- Floral scent composition was highly dynamic, with trans-geraniol comprising 72.54% of the total emission at full bloom, followed by benzyl alcohol (12.96%) and methyl salicylate (3.75%), which predominantly emanated from petals and sepals. Transcriptomic analysis and inhibition assay using pathway-specific inhibitors revealed the mevalonate (MVA) pathway as the precursor source for trans-geraniol biosynthesis. Together with elevated expression of both cytosol-localized geraniol pyrophosphate synthase EoGDPS and geraniol synthase EoTPSa1, we elucidated an unusual cytosolic biosynthesis route for geraniol in *E. oxypetalum* petals.
- Our study on *E. oxypetalum* scent emission and its biosynthesis offers a comprehensive profile of cactus floral scent profiles and presents a rare case of cytosolic geraniol biosynthesis using MVA pathway-derived substrates. These findings underline the unique metabolism of cactus flower volatiles, which hold a promise to be applied in the development of novel perfumes.

## Introduction

Angiosperm flowers emit diverse and complex fragrance of volatile organic compounds (VOCs) to attract pollinators and repel florivores (Dudareva *et al*., 2013; Dötterl & Gershenzon, 2023). The composition and emission patterns of floral VOCs exhibit considerable spatial and temporal variability (Zhang *et al*., 2022), creating distinct and characteristic fragrance. Species of Cactaceae, often cultivated as ornamental plants, emit diverse floral scents with pollinator-related chemical composition and temporal patterns. Usually, bee- or bird-pollinated cacti have scentless diurnal flowers, while nocturnal-blooming cacti emit a contrasting spectrum of volatiles. The bat-pollinated flowers usually produce sulfur-containing VOCs, causing unpleasant cabbage-like odor. Whereas moth-pollinated ornamental species usually emit complex blends of terpenoids and benzenoids volatiles during their transient nocturnal flowering, creating attractive cactus scent (Kaiser & Tollsten, 1995). *E. oxypetalum*, a popular nocturnal flowering ornamental species from Cactaceae, emits a strong and pleasant scent that last only for a few hours during nocturnal blooming. During its explosive floral VOC emission, *E. oxypetalum* starts with a slightly phenolic odor followed by a strong fruity scent. Previous studies have identified several monoterpene volatiles from moth-pollinating cacti, some of which have been widely used in fragrance industries for their fruity scent (Chen & Viljoen, 2010). However, these sporadic reports usually neglected the highly dynamic composition and biosynthetic origin of cactus scent, limiting our understanding of this highly specialized taxa.

Terpenoid volatiles contribute greatly to floral scent diversity (Abbas *et al*., 2017). Despite their great structural possibility, all terpenoid volatiles are directly derived from common iso-pentyl diphosphate precursors, which are condensed from the common C_5_ building blocks isopentyl diphosphate (IPP) and dimethylallyl diphosphate (DMAPP) (Muhlemann *et al*., 2014). Two alternative and compartmentally separated pathways synthesize IPP and DMAPP in plants. The mevalonate (MVA) pathway, located in the cytosol and peroxisome, synthesizes these C_5_ building blocks from acetyl-CoA, while the plastidic methylerythritol phosphate (MEP) pathway synthesizes IPP and DMAPP from pyruvic acid and glyceraldehyde triphosphate. It has been long accepted that these two pathways preferentially provide substrate for different classes of terpenoids (Dudareva *et al*., 2005). In general, the MVA pathway provides substrate for sesquiterpene synthesis, while monoterpene synthesis utilizes MEP-derived precursors in plastids. Recent work suggests that these two pathways are connected by species- and tissue-specific metabolic cross-talk (Vranová *et al*., 2012), providing additional possibilities for terpenoid metabolism.

The monoterpene geraniol is widely distributed in floral scent, and it has been intensively applied in cosmetic and fragrance industries for its sweet rouse-like scent. Specifically, geraniol is synthesized from geranyl pyrophosphate (GPP), the direct product of IPP and DMAPP condensation. Two different routes have been proposed for geraniol formation in previous studies. Usually, a plastid-localized geraniol synthase (GES) from the canonical terpene synthase (TPS) family (Chen *et al*., 2011) catalyzes the formation of geraniol utilizing the plastid MEP derived GPP (Dudareva *et al*., 2005) via carbocation mechanism (Christianson, 2017) (MEP-TPS case). This case was first established in snapdragon flowers (Dudareva *et al*., 2005) and confirmed later in numerous other plant species. Interestingly, a recent study in several Rosaceae species demonstrated that the cytosolic MVA derived GPP (Conart *et al*., 2023) was directly hydrolyzed by nudix hydrolase (NUDX) (Magnard *et al*., 2015) and phosphatase to form geraniol (MVA-NUDX case). Moreover, a transgenic study in *Nicotiana benthamiana* suggests that the geraniol biosynthesis potential of cytosol mainly rely on the local GPP pool and geraniol synthase abundance (Dong *et al*., 2016). These cases raise the possibility of cytosolic geraniol biosynthesis utilizing MVA-derived GPP and TPS geraniol synthase (MVA-TPS case), instead of a special nudix hydrolase. However, almost all GES characterized so far are plastid-localized (Bartram *et al*., 2006; Gutensohn *et al*., 2013; Opitz *et al*., 2014; Mendoza-Poudereux *et al*., 2015; Abbas *et al*., 2019), preventing this MVA-TPS case from validation.

In this study, we used the widely-loved ornamental cactus *E. oxypetalum* as a model system to investigate the emission dynamics and biosynthetic route of cactus floral scent. We demonstrated the temporal composition change during its transient nocturnal flowering and proved that both petal and sepal were the major source of floral VOCs emission. By combining transcriptomic and biochemical evidences, we functionally characterized a cytosol localized geraniol synthase *in vitro*. Pathway-specific inhibitor assay showed that MVA pathway mainly provided the substrate for geraniol biosynthesis, while biochemical quantification of metabolites indicated that starch was used as carbon source for the synthesis of terpenoids volatiles. Together, our results proved a rare MVA-TPS case of geraniol biosynthesis, and broadened our understanding of volatile synthesis in cactus family.

## Materials and methods

### Plant materials and chemicals

Plant materials of *E. oxypetalum* were collected from 5-years old cottage seedlings at the *E. oxypetalum* planting demonstration base of Sichuan Yuanlan Agricultural Development Co., Ltd. in Zizhong, Sichuan, China (29°40′N, 104°45′E). The seedlings were grown under semi-natural conditions with pine needle mulch soil. To aid in the development of healthy flowers, a specialized phosphate-potash-amino acid mixture was applied as a foliar fertilizer. Shade roofs were used during summer to protect plants from excessive sunlight. For RNA-seq, qRT-PCR, molecular cloning and biochemical experiments, samples from various parts of the flowers were collected in September, immediately frozen in liquid nitrogen and stored at −80L. The materials used for SPME-GC/MS were collected into glass headspace bottles, immediately frozen and stored in dry ice. Floral scent volatile samples for DHC-GC/MS were collected in October with headspace adsorption. All chemical reagents used were purchased from Shanghai Aladdin Biochemical Technology Co. Ltd. (www.aladdin-e.com).

### Floral VOCs collection and GC-MS analysis

For *in situ* VOCs collection, floral scent was sampled hourly from 19:00 p.m. to 7:00 a.m. the following day using a push-pull headspace sampling system. Briefly, precleaned (120L for 1h) polyethylene terephthalate (PET) bags (35 × 43 cm) were securely attached to the pedicel to enclose the flower. Filtered air was circulated through the bag at a flow rate of 1,000 ml min^−1^ using a battery-operated pump connected to Teflon tubing. The incoming air was purified by an activated charcoal filter to remove particles and VOCs, and a copper tube coated with potassium iodide to scrub for ozone. Air was sucked out of the bag at a flow rate of 200 ml min^−1^ through a stainless-steel adsorbent cartridge packed with 155 mg of Tenax TA, 66 mg of Carbopack B and 75 mg of Carbopack X. VOCs were collected from three flowers simultaneously, each from a different plant. After sampling, cartridges were sealed with Teflon-coated brass caps and stored at 4L until analysis. Blank measurements from empty PET bags were done to characterize impurities originating from sampling or analysis system.

VOCs trapped in adsorbent cartridges were thermally desorbed (TD100-xr, Markes International Ltd, Llantrisant, UK) at 250L for 10 min and analyzed on a gas chromatography-mass spectrometry (GC-MS) (7890A GC, 5975C VL MSD, Agilent Technologies, Santa Clara, California, USA). The VOCs were first cryofocused at −10 L and then injected into HP-5MS capillary column (30 m × 0.25 mm, film thickness 0.25 μm), using helium as carrier gas at a flow rate of 1.0 ml min^−1^ with temperature program as follows: 40L, held 1 min; raised at 5L min^−1^ to 210L, raised at 20L min^−1^ to 250L, held 8 min.

For flower organ VOCs extraction, VOCs were sampled using SPME fiber (SUPELCO 57328-U, PDMS) for 40 min at 50 L. VOCs adsorbed by SPME fibers were thermally desorbed at 230 L before analyzed on GC-MS (SHIMADZU GCMS-QP2010) with column Rtx-5. Helium was used as carrier gas at a flow rate of 1.0 ml min^−1^ using temperature program as follows: 50 L, held 3 min; raised at 6 L min^−1^ to 200 L, raised at 10 L min^−1^ to 240 L, held 6 min. Chromatograms were analyzed using PARADISe v.6.0.1 (Quintanilla-Casas *et al*., 2023). Compounds were identified using pure standards when available, or tentatively identified using NIST20 library. The VOC concentrations in blanks were subtracted from those in the samples.

### Inhibition assay for MVA and MEP pathways

In the inhibition experiment, we administered 100µM solution of the MEP-specific inhibitor fosmidomycin or the MVA-specific inhibitor mevinolin (Dudareva *et al*., 2005) by uniformly injecting 1ml of the solution into the junction of petals and perianth tubes at three specific time points: 9:00 p.m. the day before bloom, 9:00 a.m. and 7:00 p.m. The control group was injected with ddH_2_O. Floral scent samples were collected at 0:00 a.m. the next day. Throughout the experiment, the chosen inhibitors did not cause any noticeable effects on the appearance of the flowers. For MVA inhibitor assay, the lactone hydrolysis of mevinolin were performed following previous report (Kita *et al*., 1980).

### RNA-seq and analysis

Total RNAs were extracted from *E. oxypetalum* petal using an improved hexadecyl trimethyl ammonium bromide (CTAB) method. At each sampling stages: pre-bloom (9:00 a.m.), blooming (S06) and post-bloom (S13), three replicates of DNBSEQ RNA-seq libraries were prepared and quality-checked. The libraries were sequenced using the MISEQ-2000 platform by BGI, resulting in 150 bp paired-end reads. Subsequently, the raw sequencing reads underwent a quality control and cleaning process (Bolger *et al*., 2014), involving the removal of adaptors, ambiguous N-containing reads, and low-quality reads. The resulting clean data were then *de novo* assembled into transcripts using Trinity (Grabherr *et al*., 2011; Haas *et al*., 2013).

For sequence annotation, all assembled transcripts were compared to NCBI Nr, swiss-prot, GO (The Gene Ontology Consortium *et al*., 2023) and KEGG databases (Kanehisa & Goto, 2000) with an E-value threshold at 1e^−5^ for BLAST operations (Altschul *et al*., 1990). The Pfam protein database (Mistry *et al*., 2021) was searched using HMMER program (Eddy, 2011) with an E-value threshold set at 1e^−5^. Annotation integration and functional categorization by GO terms were performed by Trinotate (Duarte *et al*., 2021). Later, abundance estimation and TMM (Trimmed mean of M-values) was calculated using RSEM (Li & Dewey, 2011). Differential expression analysis was conducted using DESeq2 (Love *et al*., 2014), with a false discovery rate (FDR) threshold of < 0.01 and an absolute value of log_2_ ratio ≥2 to identify differentially expressed genes. GO term enrichment analysis was performed using GOseq R package (Young *et al*., 2010) based on the Wallenius non-central hyper-geometric distribution. Sequence clustering was carried out using MUSCLE (Edgar, 2004), ModelTest-NG (Darriba *et al*., 2020), RAxML-NG (Kozlov *et al*., 2019) and iTOL (Letunic & Bork, 2021).

### qRT-PCR for genes responsible for trans-geraniol synthesis

Specific primers for qRT-PCR were designed with NCBI. Total RNAs were extracted from *E. oxypetalum* petals with Sangon Biotech Plant Total RNA Isolation Kit (Sangon Company, https://www.sangon.com). Full-length cDNA was then reverse transcribed using Takara Prime Script™ RT reagent Kit (Perfect Real Time) (https://www.Takarabiomed.com.cn). qRT-PCR was performed with Takara RR430B TB Green Fast qPCR system, using independent RNA preparations as biological replicates. *Ubqln* gene transcripts were used as control. The relative expression levels of target genes were calculated by 2^−ΔΔCt^. The specificity of each primer pair was verified by agarose gel electrophoresis, melting curve analysis and sequence homology with local script. All relevant primer sequences were given in supplementary materials (Table S8).

### Subcellular localization of EoTPSa1 and EoGDPS

Putative sequences of *EoTPSa1* and *EoGDPS* were selected by hmmscan with Pfam models (PF01397.2, PF03936.19, PF19083.3) against the transcriptome assembly. Alignments were performed using MUSCLE (Edgar, 2004), and sequence clustering was performed with RAxML-ng. Primers were designed to amplify specific cDNA fragment with Takara PrimeSTAR® Max PCR system. We utilized SignalP 6.0 for subcellular localization predicting. Later, the transient gene expression in *N. benthamiana* leaves following established procedures with pSuper1300 and *Agrobacterium tumefaciens* strain GV3101. After infiltration, the plants were kept in the dark for 24 hours. 2.5 days after infiltration, confocal images *N. benthamiana* leaves was taken using Zeiss CELL Observer SD.

### The enzymatic activity validation of EoTPSa1

Structure prediction and molecular docking of EoTPSa1 were established with Alphafold2 (Jumper *et al*., 2021) and AutoDock (Morris *et al*., 2009) respectively. For *in vitro* enzymatic analysis, the cDNA of *EoTPSa1* was amplified and ligated into the multiple cloning sites of pET-28a expression vector utilizing Takara. In-Fusion Snap Assembly Master Mix. The constructed vector was then transformed into *Escherichia coli Rosseta-gami 2 (DE3)* cells. Positive clone incubation, protein induction, extraction and purification were all carried out under the guidance of Bacterial Protein Extraction Kit (Sangon) and Ni-NTA spin columns (Sangon) according to manufacturer’s instructions. Purified EoTPSa1 protein was examined by 12.5% (w/v) SDS-PAGE.

The enzymatic assay was conducted in 0.5ml reaction buffer (50 mM Tris–HCl, 1 mM MgCl_2_, and 0.1 mM MnCl_2_) supplemented with 100 μM of geranyl diphosphate. To the reaction buffer, 0.5μg of purified EoTPSa1 protein was added. After overlaid with 100μl hexane, the reaction mixtures were incubated at 21 L for 1 min. Reaction products were extracted by vigorously vertexing for 10s, keeping on ice for 30s, and centrifugation at 12000×g for 30s at 4 L to separate the supernatant hexane. 10μl of the dehydrated hexane supernatant was collected for GC-MS analysis. Verification and quantification of the reaction products were achieved by comparing them with the NIST17 library and authentic standards.

### Biochemical characterization of the acetyl-CoA generating metabolic pathway

The samples were collected from petals at different time points on T01-T08 (Fig. S1). The biochemical experiments were conducted using reagent kits purchased from Sangon Biotech company: reducing sugar (D799393), triglyceride (D799795), free fatty acids (D799793), and acetyl-CoA (D751001). Amylum and soluble polysaccharides were separated via centrifugation at 10000 rpm for 10 minutes after homogenization. The amylum fraction was washed with ddH_2_O, while soluble polysaccharides were precipitated using 80% ethanol. Both components were then tested with the Amylum Content Assay Kit (D799325). Absorbance in each well was measured at the required wavelength using an Agilent BioTek Synergy H1 Multimode Reader.

## Results

### Trans-geraniol dominates *E. oxyptalum* floral scent and follows the unimodal emission pattern

Unlike many other plant species that bloom gradually over an extended period, *E. oxypetalum* flowers rapidly blooms during few hours. In our experiments, the flowers typically began opening after 20:00 local time, and reached full bloom around 23:00 (Fig. 1a, 1b, Video S1). To investigate the temporal patterns of floral scent emission from *E. oxypetalum*, we collected floral VOCs at 13 different stages using dynamic headspace collection, covering the entire flowering period from 19:00 to 07:00 the next morning (Fig. 1b, 1e). In total, we identified 49 VOCs, including 26 terpenoids, 5 fatty acid derivatives and 18 benzenoids (Table S1). Among them, trans-geraniol, benzyl alcohol and methyl salicylate were the dominant VOCs, collectively constituting 90.25% of the total emission (165.45 μg/flower/h) at the peak emission stage (02:00 local time, Stage 08) (trans-geraniol, 72.54%; benzyl alcohol, 13.72%; methyl salicylate, 4.55%) (Fig. 1b, Table S2). These three VOCs were also key variables in distinguishing between blooming and non-blooming stages, as revealed by the OPLS-DA analysis (Fig. 1c).

**Fig. 1.**
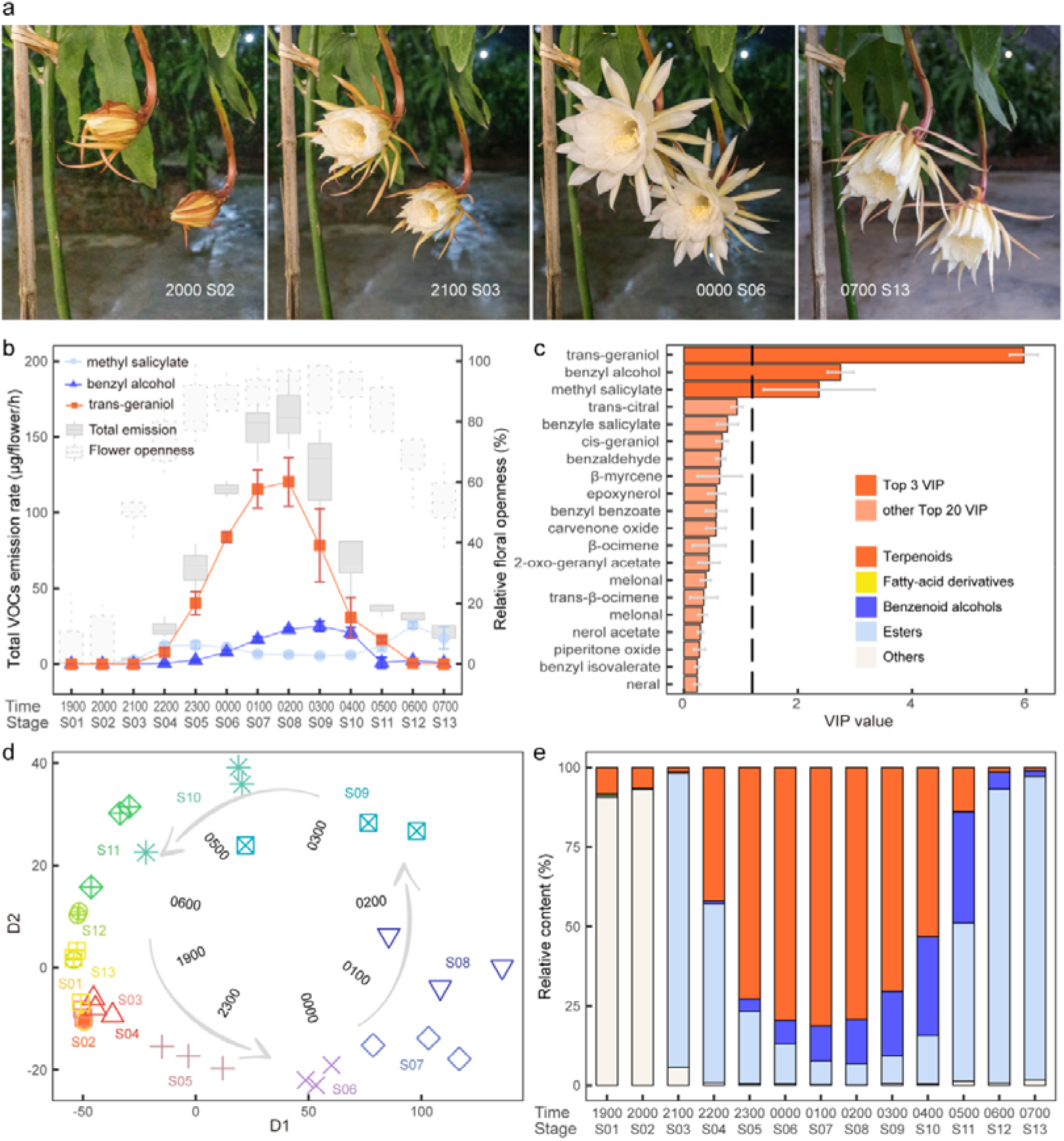
Temporal patterns of flower opening and volatile organic compounds (VOCs) emission dynamics in *Epiphyllum oxypetalum.* a: Time-lapse photos showing the rapid blooming of *E. oxypetalum* flowers. b: VOC emission rate of the categories of trans-geraniol, benzyl alcohol, methyl salicylate and total emission across all sampling stages. c: VIP value of top 20 VOCs from OPLS-DA analysis. The three major VOCs with VIP values higher than threshold were colored in brighter red. d: PCA clustering of floral VOCs content. Dots were shaped and colored according to sampling stages, time tags inside arrow rings corresponds to sampling time of nearby dots. e: Relative concentration of terpenoids, fatty-acid derivatives, benzenoid alcohols, esters and other VOCs across all sampled stages. Note that the legend is presented in c.

Total floral VOCs emission followed the unimodal pattern, in synchrony with flower openness (Fig. 1b, 1d). However, different compounds or compound classes exhibited varying temporal patterns. Initially, benzenoid esters dominated, accounting for 92.46% of the total emission shortly after flower opening at 21:00 (Stage 03). Terpenoid emission began around 20:00. (Stage 04) and became the dominant VOCs class at 01:00 (Stage 07), constituting 81.33% of the total emission. As terpenoid emission increased, benzenoid alcohols gradually surpassed benzenoid esters as the primary benzenoid VOCs, reaching their peak emission at 03:00 (Stage 09), slightly after the terpenoid peak. Subsequently, as the flowers started to wither, the emission of terpenoids and benzenoid alcohols decreased. However, from 05:00 (Stage 11), esters again constituted the majority of VOCs till the flower fully withered (Fig. 1e). Notably, trans-geraniol and benzyl alcohol displayed the unimodal emission pattern, while methyl salicylate exhibited a bimodal pattern (Fig. 1b).

### The floral scent is mainly emitted from petal and sepal of *E. oxypetalum*

To determine the primary source for floral VOCs biosynthesis and emission, we analyzed VOCs profiles of five organs, namely petal, sepal, floral tube, stamen and pistil (Fig. 2a) at three flower developmental stages: pre-bloom (S1), blooming (S8) and post-bloom (S13) using solid phase micro-extraction. The VOCs blends varied substantially depending on the flower developmental stages and the specific flower organs. The most pronounced differences among flower organs were observed at blooming, where the VOCs blends of petals and sepals were similar to each other but clearly distinguishable from those of pistils, stamens and floral tubes (Fig. 2b, Table S3). Samples other than petals and sepals from the blooming stage, as well as pistils from all stages, were clustered together. Trans-geraniol and benzyl alcohol were primarily emitted from petals and sepals, followed by pistils, floral tubes and stamens, whereas methyl salicylate was mainly emitted from pistils, followed by sepals and petals (Fig. 2c-e). Considering that petals and sepals accounted for the majority of the whole flower biomass, these organs, particularly petals, served as the primary source for trans-geraniol emission (Fig. 2f, 2g). Therefore, our subsequent analyses focused on trans-geraniol biosynthesis and subcellular compartmentalization in petal.

**Fig. 2.**
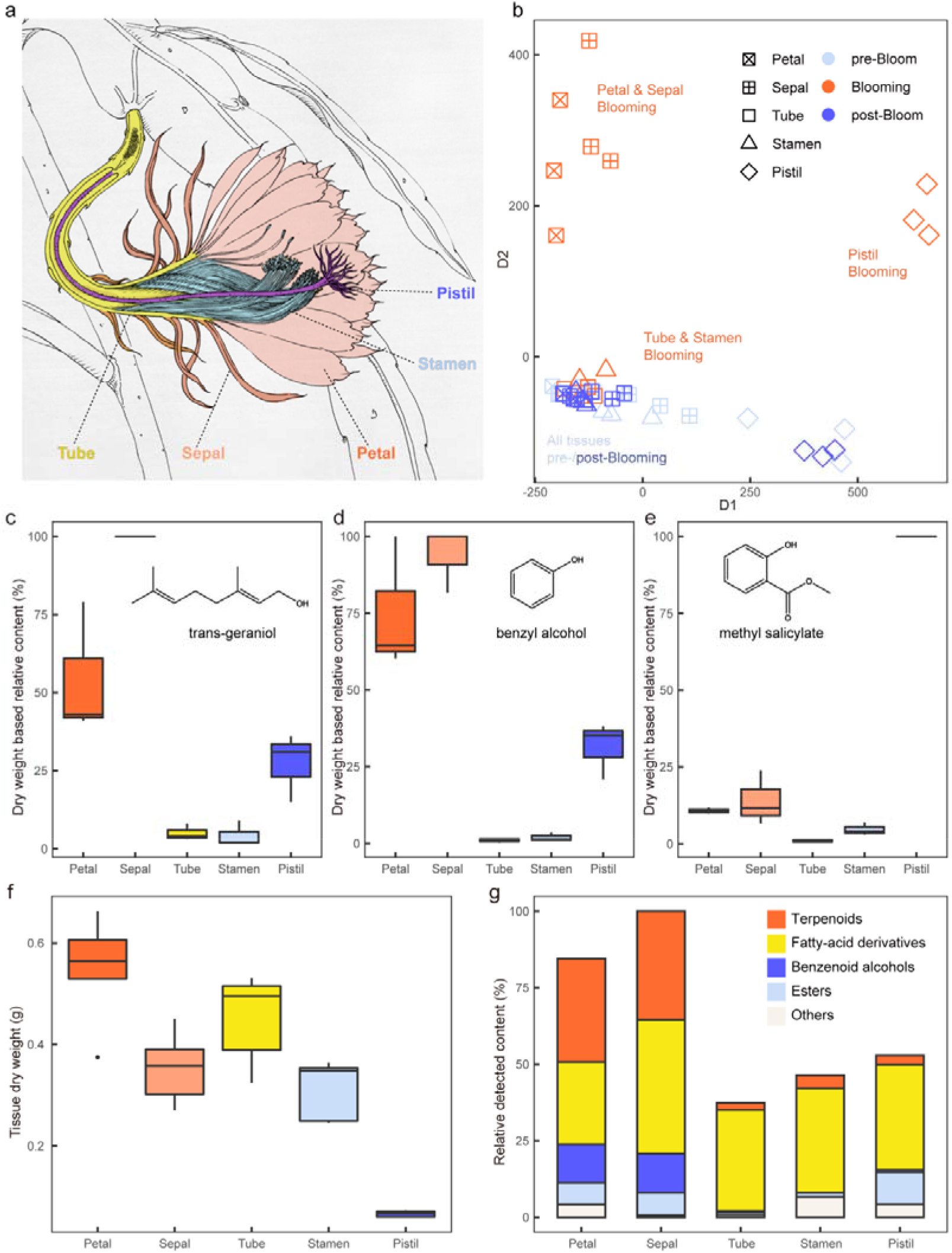
Emission patterns of five floral tissues during pre-bloom, blooming, and post-bloom stages. a: Schematic diagram of the longitudinal section of *Epiphyllum oxypetalum* floral tissues, with the five types of tissues colored differently. b: PCA clustering of volatile organic compound (VOC) content of peal, sepal, floral tube, stamen and pistil at pre-bloom, blooming and post-bloom stage. c-e: Dry weight-based content of trans-geraniol, benzyl alcohol and methyl salicylate of five floral tissues at blooming stage. f: Dry weight of five tissues sampled at blooming stage. g: Relative content of five major classes of VOCs at blooming stage, not adjusted by dry mass.

### A cytosol-localized subfamily-a terpene synthase is responsible for trans-geraniol biosynthesis

A total of 242.13 million reads were generated from nine libraries, resulting in the assembly of 164,996 unigenes and 299,340 transcripts. The average contig N50 was 2,339 bp. 64.20% of the transcripts were annotated with uniport and nr database. 45.52% of the transcripts were functionally tagged by GO database (Table S5, S6). TMM values were utilized for gene expression quantification. To identify the terpene synthases (TPS) responsible for trans-geraniol biosynthesis, we focused on transcripts annotated with Pfam TPS models (PF01397.2, PF03936.19, PF19083.3). A total of 84 unique peptides from 23 unigenes were identified as putative TPSs, with eight of them showing significant high expression levels (Fig. 3a). Sequence clustering analysis on these 23 putative TPSs from *E. oxypetalum* and 342 TPS sequences (Table S7) from 9 other species yielded 7 subfamilies, with 17 *E. oxypetalum* TPSs belonging to subfamily-a, 1 TPS to subfamily-b, 1 to subfamily-c, 1 to subfamily-e and 3 to subfamily-g (Fig. 3c). Among all the putative TPSs identified, a transcript *DN1377.2.1.2* from the subfamily-a exhibited significantly higher expression levels. This particular transcript, designated as *EoTPSa1*, exhibited approximately a 63-fold up-regulation during the blooming stage, which coincided with the peak emission of trans-geraniol.

**Fig. 3.**
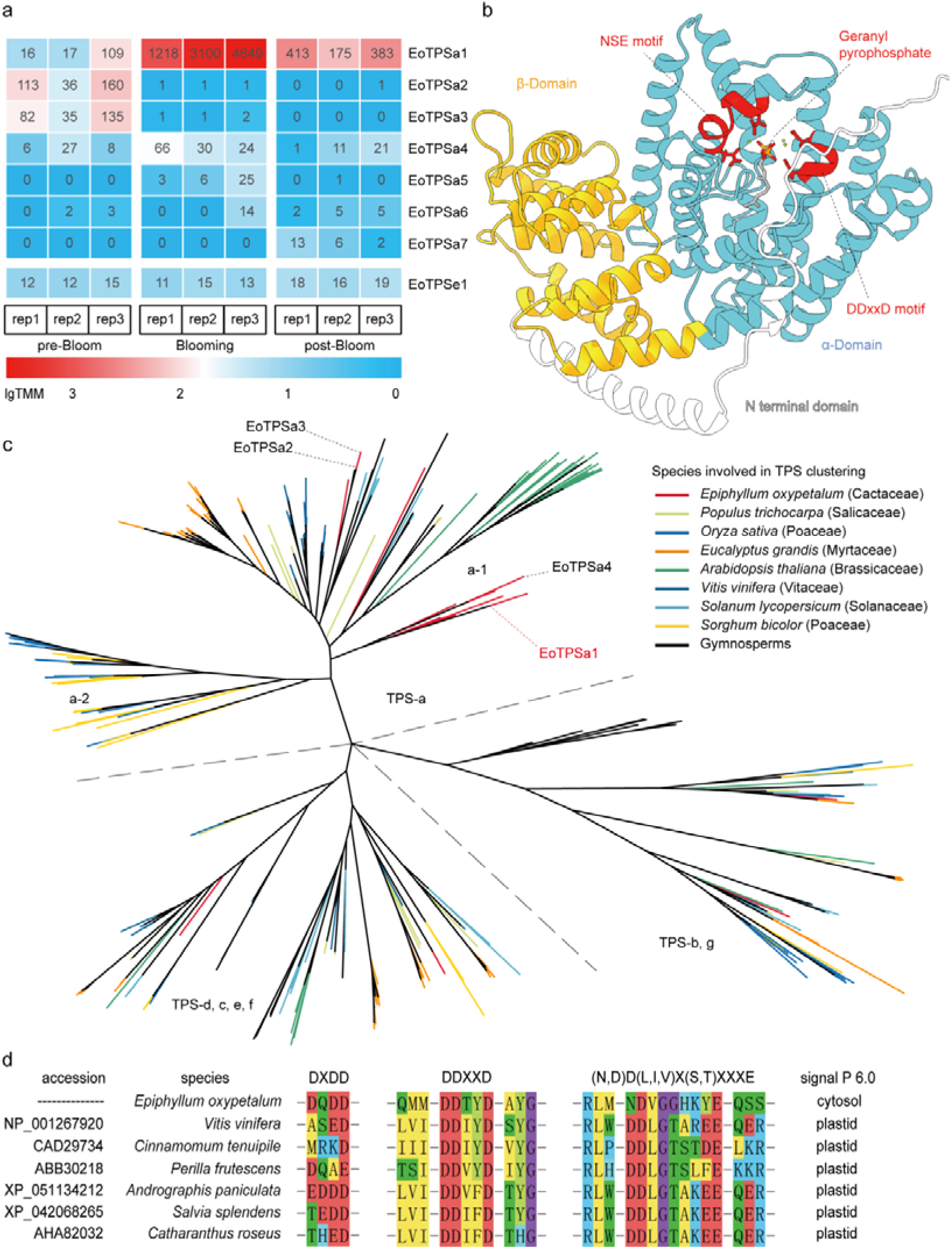
Identification of the terpene synthase for trans-geraniol biosynthesis. a: Expression patterns of eight terpene synthases identified with significant expression during pre-bloom, blooming and post-bloom stages, with each block colored according to TMM value. b: Structure prediction and molecular docking of EoTPSa1. Protein structure colored according to domain architecture and catalyzing motifs with the residues of Asp^294^, Asp^298^, Asn^440^ and Glu^448^ shown. c: Clustering of identified *Epiphyllum oxypetalum* TPSs with TPSs from other plant species. The major TPSs in *E. oxypetalum* petal marked with dashed line. d: Protein alignment of EoTPSa1 and GESs from other six species, with protein sequence around DxDD, DDxxD and NSE/DTE motifs colored according to amino acid properties. Subcellular localizations predicted with signal P 6.0 were also given.

The predicted sequence of *EoTPSa1* had an Open Reading Frame (ORF) of 1638 bp, resulting in a protein with 545 amino acids and a molecular weight of 63.29 kDa. Alignment of this protein with other functionally validated geraniol synthase (GES) revealed the presence of conserved Aspartate-rich DDxxD and NSE/DTE motifs, which were important for the magnesium-dependent prenyl diphosphate ionization in class-I mechanisms for terpene synthases. Additionally, a DxDD-like motif, commonly observed in class-II terpene synthases, was also identified (Fig. 3d). Protein structure prediction indicated the characteristic α-β domain structure of monoterpene synthases. Key amino acid residues, such as Asp^294^, Asp^298^, Asn^440^ and Glu^448^ were found to be appropriately positioned for the binding of magnesium ions (Fig. 3b). The alpha helixes in the α domain form a porous pocket that facilitates substrate binding and enables water quenching of carbon cation intermediates for trans-geraniol formation. Molecular docking simulations positioned magnesium ions and geraniol pyrophosphate accurately in the active center of the enzyme. The catalytic ability of EoTPSa1 was further verified through an *in vitro* enzymatic assay using geraniol pyrophosphate as the substrate. Notably, trans-geraniol was detected as the sole reaction product above detection limit, suggesting a high specificity of EoTPSa1. (Fig. 4).

**Fig. 4.**
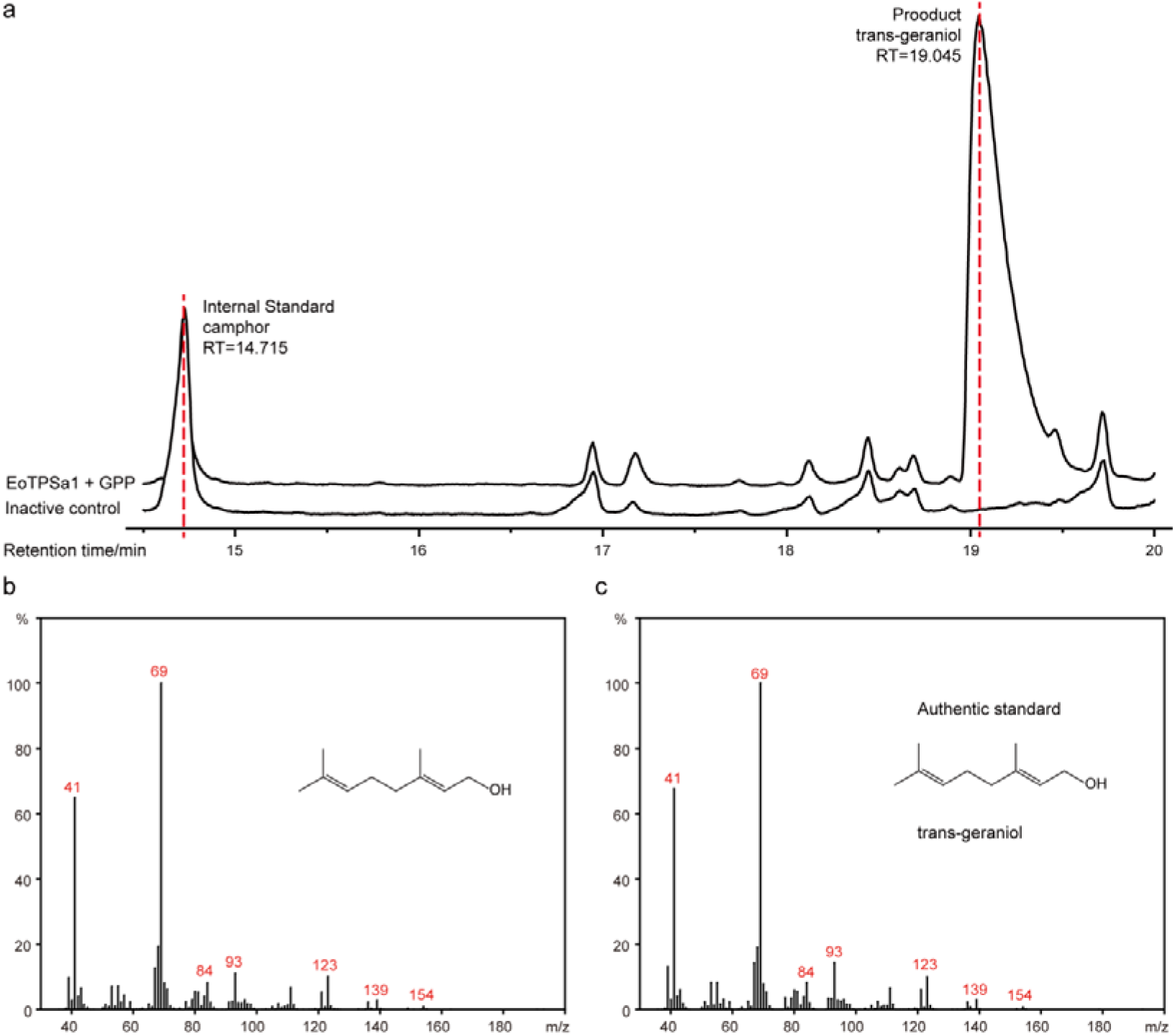
Biochemical characterization of EoTPSa1 as trans-geraniol synthase. a: TIC comparison between active enzyme and inactive control from 14.5 min to 20 min RT, showing huge trans-geraniol peak at 19.045 RT. b-c: Mass spectral fragmentation pattern of the trans-geraniol peak (RT 19.045) catalyzed with EoTPSa1 (b) and the trans-geraniol standard (c).

Additionally, EoTPSa1 was predicted to be localized in the cytosol, alongside the highly expressed geranyl pyrophosphate synthase in *E. oxypetalum*, namely EoGDPS. We further validated the cytosolic localization of EoTPSa1 and EoGDPS through *Nicotiana* transient expression system. We detected strong GFP signals in the cytosols of transformed leaf epidermis for all EoTPSa1-GFP, EoGDPS-GFP and pure GFP constructs. The GFP signal and the red chloroplast fluorescence distribution did not exhibit overlapping patterns, suggesting the absence of protein sorting activities (Fig. 5). These findings distinguish EoTPSa1 from other characterized geraniol synthases, which are typically localized in plastids.

**Fig. 5.**
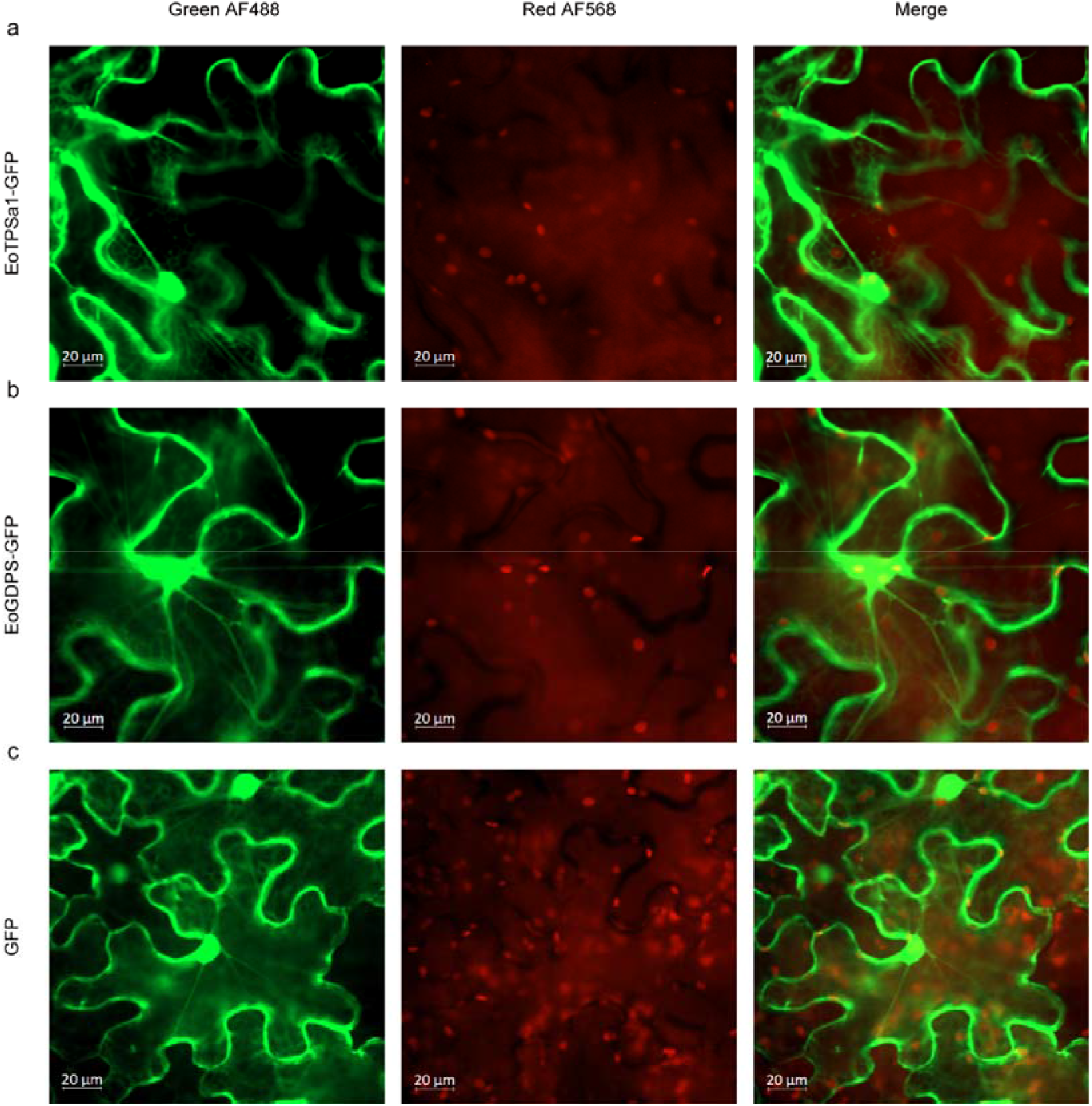
Subcellular localization of EoTPSa1 and EoGDPS. a-c: Subcellular localization of EoTPSa1-GFP, EoGDPS-GFP, and the control GFP, respectively. Confocal image was taken at AF488 green band for GFP fluorescence, and AF568 red band for chloroplast autofluorescence.

### MVA pathway mainly provides precursors for trans-geraniol biosynthesis

To validate the cytosol-localized monoterpene biosynthesis and investigate the biosynthetic origin of trans-geraniol precursors IPP and DMAPP, we examined the expression of genes involved in the MVA and MEP pathways. Interestingly, homologous genes from the MVA pathway experienced upregulation at the blooming stage, contrary to the uniformly low expression of MEP pathway genes across all three time points (Fig. 6a, Table S9). Specifically, the *HMG* gene, which encodes a putative hydroxymethylglutaryl-CoA reductase and acts as the rate-limiting enzyme of the MVA pathway, showed a 69-fold upregulation at the blooming stage compared to the pre-bloom stage. Another gene, *MVAD*, which encodes a putative diphosphate-mevalonate decarboxylase, also exhibited a two-fold upregulation. On average, during the blooming stage, the expression of the MVA pathway genes were found to be 33 times higher than that of the MEP pathway. To further investigate the predominant role of the MVA pathway in the biosynthesis of trans-geraniol, we employed pathway specific inhibitors, mevinolin for *HMG* and fosmidomycin for *DXPS*, to selectively block the MVA and MEP pathways, respectively. We then collected samples from the petal and sepal for VOCs characterization. While the VOCs of the fosmidomycin-treated flowers and untreated flowers showed similarities, there was a noticeable difference in the volatile components of the mevinolin-treated flowers (Fig. 6b, c, Table S4). This distinction was mainly driven by the emission of monoterpenes, particularly trans-geraniol, which showed 83.4% decrease in mevinolin-treated flowers but a mild increase in fosmidomycin-treated groups (Fig. 6d). In addition, mevinolin-treated flowers emitted higher amounts of ethyl salicylate but lower amounts of benzyl alcohols and methyl salicylate, whereas fosmidomycin-treated flowers emitted higher amounts of benzyl alcohol and methyl salicylate (Fig. 6e-g). This suggests that blocking either the MVA or MEP pathways may divert the carbon flow to other biosynthetic pathways, such as the benzenoid biosynthetic pathways.

**Fig. 6.**
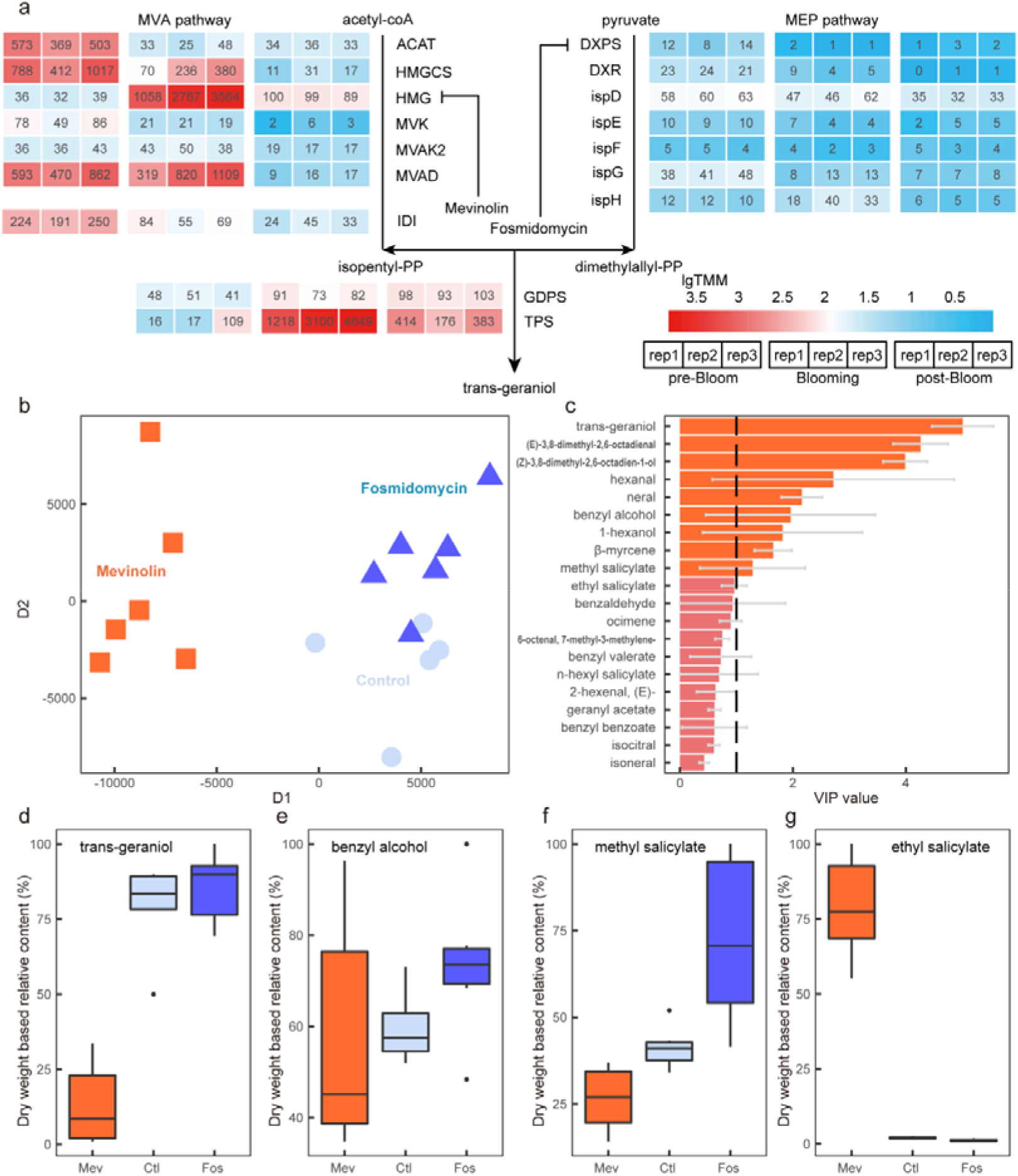
Preference of the trans-geraniol biosynthesis pathway in *Epiphyllum oxypetalum* petal. a: Expression pattern of MVA and MEP pathway genes during pre-bloom, blooming and post-bloom stages, with each block colored according to TMM value. b: PCA clustering of volatile organic compound (VOC) content of petals with mevinolin, fosmidomycin treatment group and control group at blooming. Dots were shaped and colored according to different treatments. c: VIP value of top 20 VOCs from OPLS-DA analysis. VOCs with VIP values higher than threshold were colored in orange. d-g: VOC content of trans-geraniol, benzyl alcohol, methyl salicylate and ethyl salicylate in petal of mevinolin, fosmidomycin and control group.

### Amyloplasts provide carbon source for floral scent biosynthesis

To explore the origin of the carbon source for floral VOCs biosynthesis in *E. oxypetalum*, we investigated the expression patterns of homologous genes involved in starch degradation, glycolysis, triglyceride degradation and fatty acid β-oxidation. These genes showed inconsistent expression patterns during the pre-blooming stage compared to the blooming and post-blooming stages. Notably, the beta amylase *BAM*, which catalyzes the initial step of starch degradation, showed a 6.8-fold upregulation at the full-blooming stage, while the genes involved in triglyceride degradation and fatty acid β-oxidation did not exhibit a similar trend (Fig. 7a).

**Fig. 7.**
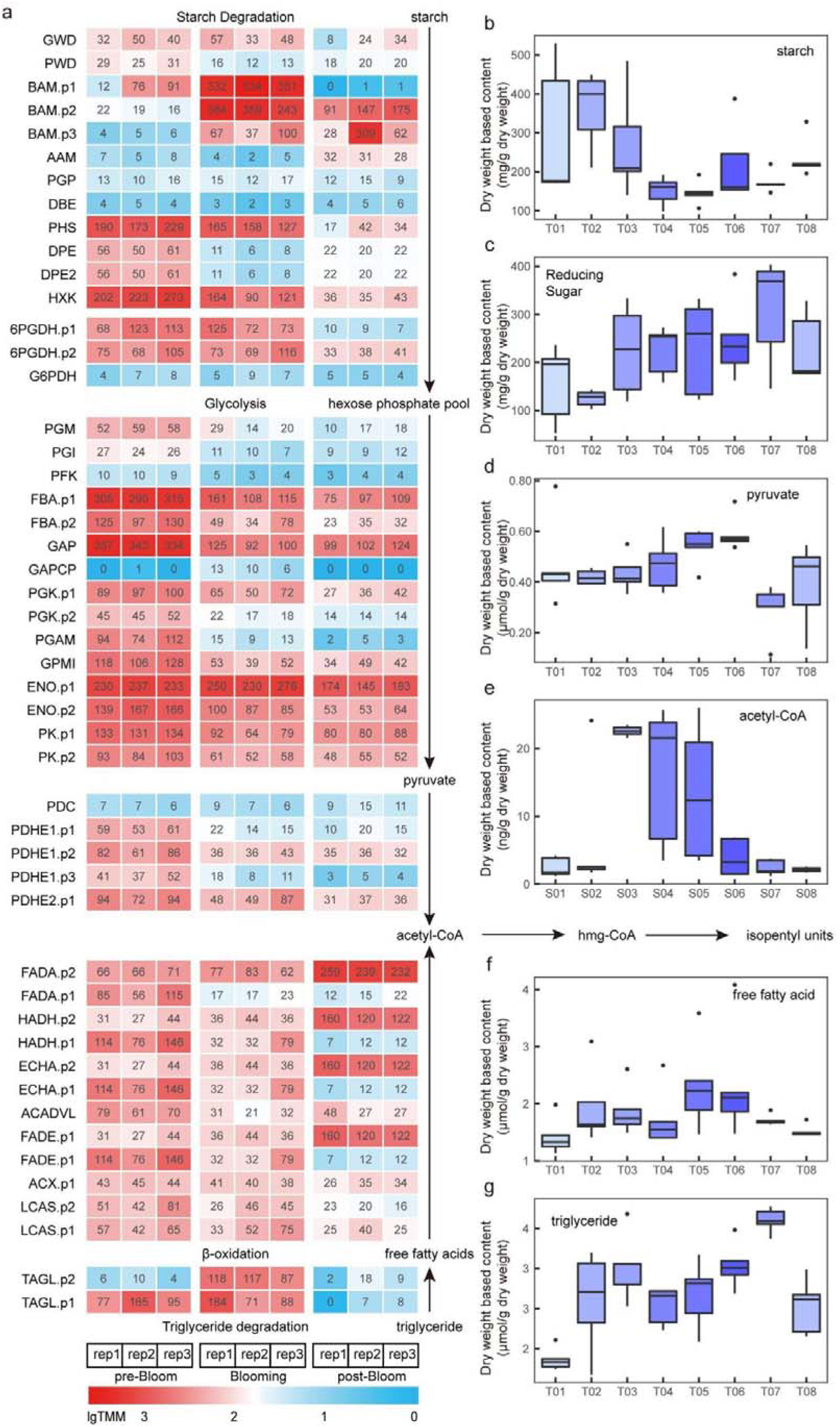
Carbon source for trans-geraniol biosynthesis in *Epiphyllum oxypetalum* petal. a: Expression pattern of starch degradation, glycolysis and β-oxidation pathway genes at pre-bloom (T03), blooming (T06) and post-bloom (T07), with each block colored according to TMM value b-g: Concentration of starch, reducing sugar, pyruvate, acetyl-CoA, free fatty acid and triglyceride in petal during T01-T08.

Later, our biochemical quantification showed that the degradation of starch began in the T02-T03 stage, leading to the accumulation of reducing sugars and a sharp increase in the content of acetyl CoA, which serves as a direct substrate for the MVA pathway. Over the next half day from T03 to T05, as metabolic activity intensified, starch degradation further increased; the concentration of reducing sugars remained consistently high, while acetyl-CoA continuously consumed. Although the total amount of pyruvate, an intermediate metabolite, was relatively low, it exhibited a significant upward trend, indicating increased physiological activity in *E. oxypetalum*. By the T05-T06 stage, due to active metabolism, starch concentration reached its lowest level, the content of pyruvate reached its peak, while acetyl-CoA was consumed to its minimum due to intense physiological processes. After the blooming stage, the acetyl-CoA content returned to its initial lowest level (Fig. 7b-e). However, not only triglycerides and free fatty acids were detected at negligible levels from T01-T07, but the trend of their changes was also independent of acetyl-CoA (Fig. 7f, 7g). Together, these findings suggest that the carbon source for trans-geraniol biosynthesis in *E. oxypetalum* originates from starch granules within the amyloplasts.

## Discussion

The biosynthesis and emission pattern of floral volatiles is highly diverse among different taxa. The Cactaceae family contains several well-known ornamental species for their unique morphological characteristics and attractive flowers (Mauseth, 2006). Extensive work has been focused on their pollination mode and ecological relevance, but the biosynthesis of their floral scent remains mostly unexplored. Here, using the famous *E. oxypetalum* as a model system, we investigated the composition and biosynthesis of its floral scent in the great detail. We first characterized its highly dynamic scent composition and then identified the petal and sepal to be the major source of VOC emission. By exploring the biosynthesis of its major component trans-geraniol, we finally showed a novel MVA-TPS case for geraniol biosynthesis in *E. oxypetalum*.

### Highly dynamic composition of *E. oxypetalum* floral scent volatiles during transient blooming

*E. oxypetalum* has impressive huge flowers and strong scent, both of which are widely distributed traits across Cactaceae. Previous investigation categorized cactus flowers into different classes according to their pollination mode, each with distinct floral VOCs composition and emission dynamics (Kaiser & Tollsten, 1995). Similar to those nocturnal blooming cacti, *E. oxypetalum* emits excessively high content of terpenoid and benzenoid volatiles during its transient blooming, which lasts for one night only. The nocturnal flowering and strong scent emission may be both due to its coevolution with insects (Guerrero *et al*., 2019) and a legacy of drought adaptation when ancient cactus first thrived in central Andeans (Hernández-Hernández *et al*., 2014). Interestingly, in our study, the composition of *E. oxypetalum* scent did not keep the same even between the short 1-hour sampling interval. Trans-geraniol was the primary compound responsible for its floral scent, and exhibited a unimodal emission pattern peaking at full blooming. The two top benzenoids in *E. oxypetalum*, benzyl alcohol and methyl salicylate, exhibited their possible substrate competition (Schuurink *et al*., 2006), as benzyl alcohol peaked around full-bloom while methyl salicylate peaked before and after full-bloom. The biosynthesis of these VOCs primarily occurred in the petal and sepal tissue, both with high surface-to-volume ratio to promote the overall scent emission. Also, trans-geraniol, benzyl alcohol and methyl salicylate are all known to convey chemical signals between plants and insects, especially trans-geraniol for hawk-moth (Foster And & Harris, 1997), and thus they may have played crucial roles in the interaction between *E. oxypetalum* and pollinators. Whether these compounds, either individually or in specific ratios, during flowering convey specific chemical messages to pollinators remains to be investigated (Raguso, 2008).

### Geraniol biosynthesis in MVA-TPS pathway

Generally, monoterpenes volatiles are synthesized with plastid-localized monoterpene synthase utilizing MEP-derived GPP (MEP-TPS case) (Dudareva *et al*., 2005). Interestingly, recent work in rose proved a special nudix hydrolase that catalyzed the hydrolysis of GPP into geraniol (MVA-NUDX case) (Magnard *et al*., 2015; Conart *et al*., 2023); However, apart from these two canonical cases, some recent reports provided several exceptions. Several cytosolic-localized geraniol synthases were characterized in *Fragaria ananassa* (Aharoni *et al*., 2004), *Lippia dulcis* (Dong *et al*., 2013), *N. benthamiana* (Wu *et al*., 2006; Dong *et al*., 2016) and *Solanum lycopersicum* (Davidovich-Rikanati *et al*., 2008; Zhou & Pichersky, 2020). Moreover, isotope labeling experiments proved MEP-dependence of sesquiterpene formation in snapdragon, along with MVA-dependent linalool formation in raspberry (Hampel *et al*., 2007). These results suggest a possible new case for monoterpene biosynthesis, in which a cytosolic TPS might utilize MVA-derived GPP for monoterpene production (MVA-TPS case). This MVA-TPS case might be more ubiquitous than the special MVA-NUDX case for geraniol biosynthesis, as a nudix hydrolase capable of hydrolyzing GPP into geraniol was only characterized for rose hybrids and *Pelargonium graveolens* (Bergman *et al*., 2021) so far, while both TPS and MVA pathway are widely distributed in higher plants.

In the current study, we have provided both biochemical and transcriptomic evidence that a cytosolic-localized TPS utilizes MVA-derived GPP to synthesize trans-geraniol in *E. oxypetalum* flowers. The cytosol localization of geraniol synthase EoTPSa1 and significantly reduced terpenoid content during mevinolin treatment provided the solid evidence for the MVA-TPS case. The special geraniol synthase EoTPSa1, which contributed most of the total scent emission at full bloom, was classified into subfamily-a during sequence clustering. Previous study has suggested sesquiterpene preference for subfamily-a TPSs (Chen *et al*., 2011), and the characterized geraniol synthases so far has diverse phylogenetic preference. This observation concurs with previous knowledge about active gene neo-functionalization during terpenoid evolution, and preference for utilizing existing enzymes for specialized metabolism with catalytic promiscuity (Weng, 2014). Incorporation of cytosolic MVA pathways rather than plastid originated substrate coupled downstream transportation can improve catalysis efficiency significantly (Dong *et al*., 2016). This special MVA pathway incorporation might be due to the elevated HMG expression at full bloom, which might share the common regulation scheme with eoTPSa1.

In our transcriptome assembly, *EoTPSa1* was the only terpene synthase encoding gene that exhibited significant expression. *In vitro* enzymatic assay identified EoTPSa1 with significant trans-geraniol biosynthesis activity. Interestingly, both isomers trans-geraniol and cis-geraniol, along with their respective oxidized products trans- and cis-citral were detected in floral scent. While the existence of citral suggests the possible dehydrogenase activity (Zhao *et al*., 2023), cis-geraniol formation indicates an uncommon lack of stereoselectivity during GPP biosynthesis. Usually, during the nucleophilic attack of IPP on the isoprene carbocation generated by the dissociation of pyrophosphate in DMAPP, the proton abstracted by the newly formed tertiary carbocation often generates an trans-structured double bond (Walsh & Tang, 2017). The formation of less common cis-structured double bonds in isopentenyl chain extension is only sometimes observed in bactoprenols formation (Walsh & Wencewicz, 2016). In order to form a cis-structured double bound, the basic group that catalyzes the departure of hydrogen ion must be positioned on the opposite side of the reaction site, which is evident by the difference between trans- and cis-stereoselective isopentyl diphosphate synthase (Guo *et al*., 2004). However, our results showed that only one GPP synthase might be active during *E. oxypetalum* trans- and cis-GPP synthesis, suggesting that the GPPS identified might be capable to form both trans- and cis-structured double bound simultaneously, and the geraniol synthase to hydrolyze these two isomers at the same time.

### Starch as primary carbon source for the synthesis of *E. oxypetalum* floral VOCs

MVA pathway utilizes acetyl-CoA as substrate, which can derive from both starch and lipid degradation. Transcriptome assembly and quantifications identified the active expression at blooming stage for both β-amylase (BAM) and TAGL, the initial steps for these two degradation pathways. Biochemical quantification revealed the existence of excessively high content of starch (up to 30% dry weight) and reducing sugars in petal, but almost complete lack of triglyceride and free fatty acids, indicating that starch may act as the primary carbon source. The drop in starch content suggests that starch degradation initiated in the morning on the day of blooming, creating hoarded acetyl-CoA for active metabolism during flower opening and volatile biosynthesis. Thus, our findings suggest that an adequate supply of substrate, the strict regulation of enzymes, and the selection of the cytoplasm as the site for geraniol synthesis collectively constitute the mechanism that allows *E. oxypetalum* to release a substantial number of volatile compounds in such a short flowering period, and distinguishing it from other plants in the cactus family with usually longer flowering length and less fragrance emission.

## Acknowledgements

This work was supported by the National Nature Science Foundation of China (32271692, 32100326), and the Department of Science and Technology of Sichuan Province of China (2022ZHXC0009, 2023YFSY0054). We further wish to thank Yudong Ma, Wei Wu and Jin Dai for their assistance in sample collection, and Bolei Jiao, Xinrui Liu and Chuanfang Wu for sharing the *N. benthamiana* plants and pSuper1300, pET-28a plastids. We also would like to thank Qijun Deng and Wei Ni from Sichuan Bowu Cultural Development Co. Ltd. for the drawing of *E. oxypetalum* flowers.

## Competing interests

None declared

## Author contributions

Yiyang Zhang, Yuhan Zhang, LT and QS designed the experiments. The top seven authors and WT jointly completed the collection and preparation of samples. Yiyang Zhang and Yuhan Zhang quantified VOCs content with GCMS. YiYang Zhang analyzed GCMS and transcriptome data. Yuhan Zhang and QT performed molecular cloning, enzymatic assay and localization studies. LW performed biochemical quantifications. Yiyang Zhang plotted the figures and wrote the first draft. Yuhan Zhang, QS, TL, HX, QT, ML and ZL revised the manuscript. Yiyang Zhang and Yuhan Zhang contributed equally to this work.

## Data Availability

All data needed to evaluate the conclusions in the paper are present inthe paper and/or the Supplementary Materials. The transcriptome raw reads for 9 individuals in this study have been deposited in the National Genomics Data Center (https://ngdc.cncb.ac.cn) under accession number PRJCA024038.

## Supporting information

Table S1. Content matrix of compounds detected in *Epiphyllum oxypetalum* floral scent.

Table S2. Absolute content of top eight volatiles detected in floral scent.

Table S3. Content matrix of compounds detected in samples of 5 floral tissues at pre-bloom, blooming and post-bloom.

Table S4. Content matrix of compounds detected in pathway inhibitor assay.

Table S5. Transcriptome expression matrix of samples at pre-bloom, blooming and post-bloom.

Table S6. Transcriptome annotation matrix of samples at pre-bloom, blooming and post-bloom.

Table S7. Protein sequences used for TPS family sequence clustering.

Table S8. Primers used in this study.

Table S9. Results matrix for qRT-PCR measurement of genes involved in MVA, MEP and geraniol synthesis.

Figure S1. Sampling time for photography, transcriptome, headspace and biochemical experiments.

Video S1. Time-lapse video for the nocturnal blooming of *Epiphyllum oxypetalum*

## Notes

### Competing Interest Statement

The authors have declared no competing interest.

### Summary of Updates

author affiliations updated; Supplemental files updated;etc.

