## Supplementary figures and images for "Floral scent emission of *Epiphyllum oxypetalum*: identification of a novel cytosol-localized geraniol biosynthesis pathway"

### Fig. S1

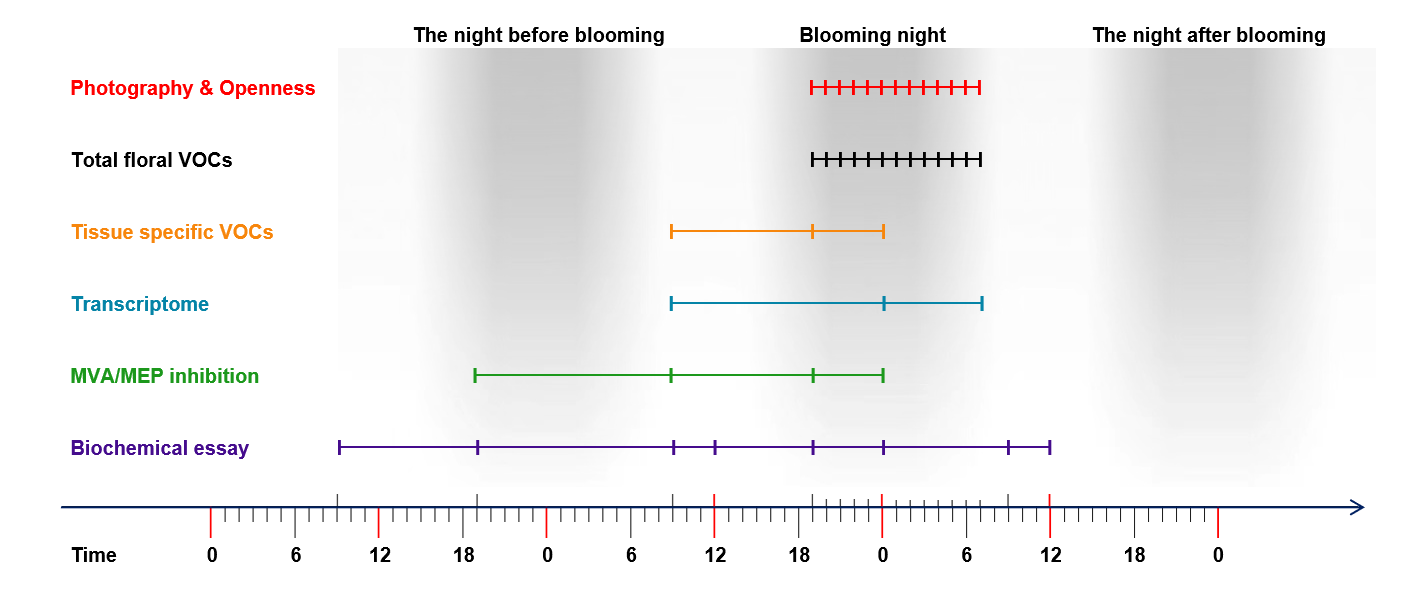
